# Re-routing metabolism by the mitochondrial pyruvate carrier inhibitor MSDC-0160 attenuates neurodegeneration in a rat model of Parkinson’s disease

**DOI:** 10.1101/2022.01.17.476616

**Authors:** David Mallet, Raphael Goutaudier, Emmanuel L. Barbier, Sebastien Carnicella, Jerry R. Colca, Florence Fauvelle, Sabrina Boulet

## Abstract

**Background:** A growing body of evidence supports the idea that mitochondrial dysfunction might represent a key feature of Parkinson’s disease (PD). Central regulators of energy production, mitochondria are also involved in several other essential functions such as cell death pathways and neuroinflammation which make them a potential therapeutic target for PD management. Interestingly, recent studies related to PD have reported a neuroprotective effect of targeting mitochondrial pyruvate carrier (MPC) by the insulin sensitizer MSDC-0160. As the sole point of entry of pyruvate into the mitochondrial matrix, MPC plays a crucial role in energetic metabolism which is impacted in PD. This study therefore aimed at providing insights into the mechanisms underlying the neuroprotective effect of MSDC-0160.

**Methods:** We investigated behavioral, cellular and metabolic impact of chronic MSDC-0160 treatment in unilateral 6-OHDA PD rats. We evaluated mitochondrial related processes through the expression of pivotal mitochondrial enzymes in dorsal striatal biopsies and the level of metabolites in serum samples using nuclear magnetic resonance spectroscopy (NMR)-based metabolomics.

**Results:** MSDC-0160 treatment in unilateral 6-OHDA rats improved motor behavior, decreased dopaminergic denervation and reduced mTOR activity and neuroinflammation. Concomitantly, MSDC-0160 administration strongly modified energy metabolism as revealed by increased ketogenesis, beta oxidation and glutamate oxidation to satisfy energy needs and maintain energy homeostasis.

**Conclusion:** MSDC-0160 exerts its neuroprotective effect through reorganization of multiple pathways connected to energy metabolism.

## Introduction

Parkinson’s disease (PD), mainly characterized by the progressive loss of dopaminergic neurons of the substantia nigra pars compacta (SNc), is the second neurodegenerative pathology worldwide. It remains incurable, possibly because of its particularly complex etiology[1]. Various mitochondrial alterations have been highlighted as major actors in the cascade of events leading to the degeneration of dopaminergic neurons[2–5]. Indeed, beyond their prominent role in energy metabolism, mitochondria are involved in several other essential functions dysregulated in PD, including the activation of cell death pathways such as the mammalian target of rapamycin (mTOR), which regulates autophagy [3, 6, 7]. Alteration of mitochondria has also been shown to be associated with neuroinflammatory processes, especially through the accumulation of the inducible isoform of nitric oxide synthase (iNOS), which leads to nitric oxide formation, an inhibitor of mitochondrial respiration[8, 9]. Yet most of the therapeutic strategies focusing on these processes have failed, probably because treatments have targeted them individually[10–12]. In contrast, reprogramming mitochondrial metabolism, by targeting upstream reactions in the signaling cascades, may represent a more efficient therapeutic strategy in PD[13, 14].

Among various potential sites of action, pyruvate metabolism, critical for energy generation[15], has emerged as a promising therapeutic target. Indeed, as the end-product of glycolysis and at the entry to mitochondrial metabolism, pyruvate represents the main fuel supply of the tricarboxylic acid (TCA) cycle[15] and its metabolism plays a key role in cell homeostasis[16]. Strikingly, abnormally high levels of pyruvate have been observed in PD patients[15, 17, 18], reflecting alteration in its metabolism. Consistently, using nuclear magnetic resonance (NMR)-based metabolomics in a recent study combining three different PD animal models and two independent PD patients cohorts, we have identified a set of metabolic dysregulations which strongly suggest that modification of pyruvate metabolism is associated with PD pathophysiology and progression[19]. In particular, glycolytic metabolites, including pyruvate, were increased whereas the TCA cycle metabolites remained stable. This suggests that cytosolic glycolysis and mitochondrial TCA cycle are decoupled, impacting the supply of energy. Concomitantly, ketone bodies and amino acids were increased, suggesting that other fuel sources were being used to maintain the TCA cycle[19].

Coordination between glycolysis and mitochondrial activities depends on the entry of pyruvate into the mitochondrial matrix, by the mitochondrial pyruvate carrier (MPC) which thus appears pivotal for modulating the cell energy production[20–22]. Indeed, previous studies have shown that MPC knockdown results in compensatory use of other substrates, such as amino acids, fatty acids and ketone bodies, a phenomenon highly reminiscent of our previous work[23, 24]. Additionally, MPC inhibition by an insulin sensitizer thiazolidinedione compound (TZD) compound, the MSDC-0160, in a MPTP mice model of PD has been found to limit dopaminergic denervation and to reduce associated motor impairments, through a mechanism that involves modulation of mTOR signaling. These effects of MSDC-0160 included normalization of autophagy, decreased neuroinflammation and reduced iNOS expression[25]. Interestingly, TZD such as MSDC-0160, which specifically target MPC[26], can reduce pyruvate entry into mitochondria. Reduced activity of MPC in cultured dopaminergic neurons protects them from excitotoxic death[27]and observational clinical studies have shown that, compared to other antidiabetic drugs, the use of TZDs in diabetic patients is associated with a reduction of PD incidence [28].

In light of these observations, we hypothesized that MPC dysfunction may be the cause of the metabolic dysregulations we observed in our previous study[19], and that inhibiting MPC could therefore be neuroprotective. To validate the neuroprotective action of MSDC-0160, we first characterized its effects at a behavioral and histological level in unilateral 6-hydroxydopamine (6-OHDA) PD rats. Then we explored the metabolic cell processes underlying the mechanism of action of MSDC-0160 in this model, initially focusing on mitochondrial metabolism known to be altered by 6-OHDA cytotoxicity [29–31].

As predicted from murine models of PD, behavioral and histological results confirmed that MSDC-0160 treatment protected against dopaminergic denervation and resulting motor dysfunction in the 6-OHDA rat model. Exploration of the cell processes involved demonstrated that treatment with MSDC-0160 normalized mTOR signaling and iNOS expression in these PD animals. Finally, from a metabolic standpoint, we found that MSDC-0160 treatment was associated with an increased use of lipids and ketones bodies as alternative fuel sources. Altogether, these findings suggest that MSDC-0160 treatment could exert its neuroprotective action through two different ways: the promotion of compensatory metabolic adaptation and the limitation of neuroinflammatory and apoptotic mechanisms. Altogether, these results strongly support MPC inhibition as a highly relevant therapeutic strategy in PD by counteracting some quintessential pathophysiological processes.

## Material and Methods

### • Animals

Experiments were performed on adult male Sprague-Dawley rats (Janvier, Le Genest-Saint-Isle, France), weighing approximately 250 g (5 weeks old) at the beginning of the experiment. They were housed under standard laboratory conditions with reversed light-dark cycle (12 h/light/dark cycle, with lights ON at 7 p.m.) and with food and water available *ad libitum*.

### • 6-OHDA rat model

As previously described[32, 33], rats were subcutaneously injected with desipramine (15 mg/kg) 30 min before 6-OHDA (Sigma-Aldrich, Saint Quentin-Fallavier, France) injection in order to protect the noradrenergic neurons. All animals were then anesthetized by intraperitoneal injection of Ketamine (Chlorkétam, 100 mg/kg, Mérial SAS, Lyon, France) and Xylazine (Rompun, 7 mg/kg, Bayer Santé, Puteaux, France) and placed in a stereotactic frame (Kopf instrument, Phymep, Paris, France). The stereotaxic coordinates of the injection sites, according to the stereotaxic atlas of Paxinos and Watson[34] and relative to interaural line, were as follows: incisor bar placed at −3.2 mm; anteroposterior (AP) = +3.8 mm / lateral (L) = +/− 2.2 mm / dorsoventral (V) = −8.1 mm. Animals received a unilateral injection (counterbalanced side) of 3 μl 6-OHDA (3 μg/μl, 6-OHDA group) or 0.9% NaCl (sham group), at a flow rate of 0.5 μl/min.

### • MSDC-0160 treatment

In accordance with previous studies[25, 35], rats received a daily dose of MSDC-0160 (30 mg/kg, Metabolic Solutions Development Company, Kalamazoo, USA) or placebo (1% methylcellulose with 0.01% Tween 80) by oral gavage, 4 days before unilateral injection of 6-OHDA or NaCl into the SNc (Sham placebo n=12; Sham MSDC-0160 n=9; 6-OHDA placebo n=13; 6-OHDA MSDC-0160 n=14), and continuing for 14 days after surgery, during the development and stabilization of the 6-OHDA lesion [36]. Finally, rats were euthanized 10 h after the last administration of MSDC-0160 and after tail-blood collection.

### • Behavioral assessment

At the end of treatment, locomotor asymmetry was measured with the cylinder test[37]. Animals were placed for 5 min in a cylinder, 20 cm high and 20 cm in diameter, and their forelimb use during rearing was videotaped (OBS studio open-source software). The test was carried out blind to the experimental conditions. Slow motion recorded videos were viewed to count the number of forelimbs uses, defined by the placement of the whole palm on the wall of the arena. Unilateral lesion with 6-OHDA resulted in preferential use of the ipsilateral forelimb paw [37].

### • Histological analysis

After behavioral assessment (i.e. 2 weeks after 6-OHDA or saline injection) and blood collection, rats a light gaseous anesthesia were euthanized by decapitation and brains were immediately frozen in liquid nitrogen and stored at −80 °C.

They were then cut in 14 μm coronal sections at −20 °C using a cryostat (Microm HM 525; Microm, Francheville, France).

Only animals with correct cannula implantation, verified by Cresyl violet staining, were included in the study to ensure homogeneous induction of the lesional process in every experimental condition (Additional file 1: Fig. S1).

Tyrosine hydroxylase (TH) immunostaining was carried out as previously described[19, 32, 33] on striatal and SNc sections at levels of interest (striatum: AP: +2.0/+1.6/+1.0; SNc AP: −4.8/−5.2/−5.6 related to Bregma). Briefly, after post-fixation with 4% paraformaldehyde, slices were incubated with an anti-TH antibody (mouse monoclonal MAB52 80, Millipore, France, 1: 2500) overnight at 4°C. Then slices were incubated with biotinylated goat anti-mouse IgG antibody (BA-9200, Vector Laboratories, Burlingame, CA, USA; 1: 500) and immunoreactivity revealed with Avidin-peroxidase conjugate (Vectastain ABC Elite, Vector Laboratories Burlingame, CA, USA).

Quantification of the extent of striatal dopaminergic denervation was determined with ICS FrameWork computerized image analysis system (Calopix, 2.9.2 version, TRIBVN, Châtillon, France) coupled with a light microscope (Nikon, Eclipse 80i). After drawing masks from three striatal levels, optical densities (OD) were measured. OD are expressed as percentages relative to the mean optical density obtained from the corresponding contralateral region for each animal.

For western blot analysis for each animal, thick sections (80 μm) of dissected dorsal striatum (DS) tissue were pooled in an Eppendorf^®^ and kept at −80°C until analysis.

### • Western blot analysis

Striatum samples were homogenized by sonication in SDS buffer (10%). After determining the dynamic range of detection for each antibody, 20 μg of total protein were resolved in 4-20% SDS-PAGE and then transferred onto polyvinylidene difluoride (PVDF) membranes using the Trans-Blot Turbo transfer system (Bio-rad). Briefly, membranes were incubated overnight at 4°C with primary antibodies of interest anti-: TH (Cell Signaling; s58844), mTOR (Cell Signaling; s2983), p-mTOR (Cell Signaling; s5536), iNOS (Cell Signaling; s13120), MPC1 (Cell Signaling; s14462), pyruvate dehydrogenase (Cell Signaling; s3205), p-pyruvate dehydrogenase (Cell Signaling; s31866), acetyl-coenzyme A acetyltransferase 1 (Cell Signaling; s44276), carnitine palmitoyltransferase 1a (Cell Signaling; s97361), glutamate dehydrogenase (Cell Signaling; s12793), vinculin (Cell Signaling; s13901), βactin (Cell Signaling; s4970). After washing, membranes were incubated 1h at room temperature with HRP-linked goat anti-rabbit antibody (Cell Signaling; s7074). Bands were detected using enhanced chemiluminescence (ECL) and their densities quantified using ImageLab software (Bio-rad). Each band was delimited manually, and densities were normalized by dividing by the density of controls[38].

### • NMR experiments

Blood was collected from the caudal vein under gas anesthesia with isoflurane (2%) and after 2 h of fasting, as previously described [19]. It was then stored on ice before rapid centrifugation at 1600g for 15 min at 4° C. The supernatant serum was removed and stored at −80°C until the day of NMR. The time before freezing never exceeded 30 min[39].

NMR tubes were filled with 60 *μ*l serum sample and 120 *μ*l phosphate buffer saline (PBS) 0.1M in D_2_O (50% of D_2_O, pH = 7.4). ^1^H NMR experiments were performed on a Bruker Advance III NMR spectrometer at 950 MHz (IBS, Grenoble, France) using a cryo-probe with a 3 mm tube holder. For each sample, two one-dimensional ^1^H-NMR spectra were acquired: one metabolite-edited spectrum using the Carr–Purcell–Meiboom–Gill (CPMG) pulse sequence and one lipid-edited spectrum using diffusion filtering (ledbpgppr2s pulse sequence). The residual water signal was pre-saturated for 2 seconds. Assignment of peaks was performed using 2-dimensional experiments and databases[40]. Then, as previously described, metabolite relative concentrations were extracted from CMPG spectra[19] and mean chain length and unsaturation were calculated from diffusion-edited spectra[41]. The total acquisition time lasted 20 minutes.

The free induction decays were Fourier transformed and manually phased with the Bruker software Topspin version 3.6.2. Then, further pre-processing steps (baseline correction, alignment, bucketing) were performed using NMRProcFlow v1.4 online (http://nmrprocflow.org). The spectra were segmented in 0.001 ppm buckets between 0 and 8.5 ppm with exclusion of residual water peaks, macromolecule signals (for CPMG) and other regions corresponding to pollution. Each bucket was normalized per spectrum to the sum of all buckets.

### • Statistical analysis

All univariate analyses were performed using Graphpad Prism 8 software (San Diego–USA). All results were expressed as mean values ± standard error of mean (SEM). Parametric analyses were performed after verification of the assumptions of normality (Shapiro-Wilk and Kolmogorov-Smirnov tests) and sphericity (Bartlett’s test). For all results, two-way ANOVA followed by a post-hoc Tukey test with correction for multiple comparisons were performed. Significance for p values was set at *α* = 0.05.

## Results

### Treatment with MSDC-0160 induced neuroprotection in unilateral 6-OHDA PD rats

We first tested whether chronic MSDC-0160 administration could protect SNc DA neurons and limit associated motor impairment in a 6-OHDA rat model with severe unilateral DA lesion.

As expected, a significant loss of dopaminergic neurons of the SNc and of their projections in the striatum was observed in 6-OHDA rats 2 weeks after surgery, (Fig. 1A and 1C; main effect of lesion: F_1,44_ = 258.7, p < 0.001), as visualized by loss of TH expression in the striatum (Fig. 1B and 1D; main effect of lesion: F_1,44_ = 37.29, p < 0.0001). The 6-OHDA-induced neuronal damage and striatal DA denervation were significantly attenuated in animals treated with MSDC-0160 (Fig. 1A and 1C; main effect of lesion and treatment and significant interaction between both factors: F_S_ > 13.62, p_S_ < 0.0004; Fig. 1B and 1D; main effect of lesion and treatment F_S_ > 8.59, p_S_ < 0.001), whereas MSDC-0160 alone had no effect on these parameters (Fig. 1A and 1C; post-hoc test of Tukey adjusted p value > 0.99; Fig. 1B and 1D; post-hoc test of Tukey adjusted p value > 0.61).

**Fig. 1:**
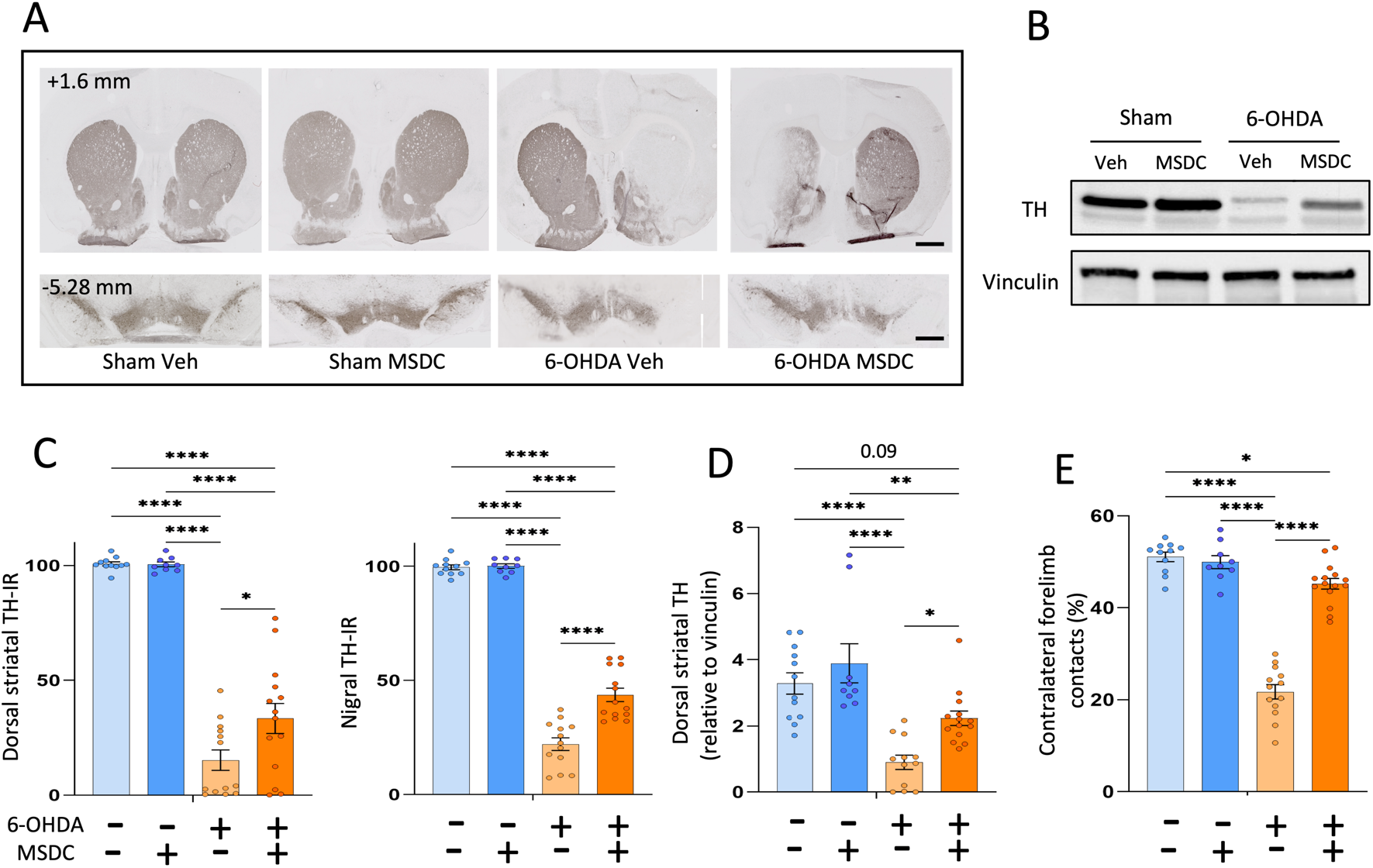
Treatment with MSDC-0160 induced neuroprotection in unilateral 6-OHDA PD rats. (A) Example of representative immunohistochemistry for TH in dorsal striatum (upper panel) and substantia nigra pars compacta (lower panel) in sham and 6-OHDA rats, with or without MSDC-0160 (AP: +1.6 mm and −5.28 mm relative to bregma). Scale bars represent 2 mm. (B) Representative western blots illustrating the level of TH in the dorsal striatum of sham and 6-OHDA rats, treated or not with MSDC. (C) Quantification of TH-IR staining loss at the dorso-striatal (left bar graph) and the nigral (right bar graph) levels, expressed as a percentage of the mean value obtained for the contralateral side. (D) Bar graph showing mean density of western blot TH/vinculin ratio in the dorsal striatum. (E) Bar graph showing contralateral forelimb contacts in the cylinder test expressed as a percentage of ipsilateral forelimb contacts. Mean ± SEM, Two-way ANOVA followed by Tukey’s post-hoc test and correction for multiple comparisons *: p ≤ 0.05, **: p ≤ 0.01, ****: p ≤ 0.0001

At the end of treatment, the evaluation of the sensori-motor function of the rats using the cylinder test showed that unilateral 6-OHDA rats had significant motor impairment, with a dramatic decrease of contralateral forelimb use consistently with previous reports[42, 43] (Fig. 1E; main effect of lesion: F_1,44_ = 161.2, p < 0.001). Chronic MSDC-0160 treatment almost completely prevented these deficits (Fig. 1E; main effect of treatment and significant interaction between lesion and treatment F_S_ > 69.46, p_S_ < 0.001).

### Chronic MSDC-0160 treatment normalized mTOR activity and iNOS expression

Previous studies have implicated the mTOR pathway in 6-OHDA-induced neurotoxicity[44], and MSDC-0160 has been shown to modify its activity [25]. We therefore measured the level of mTOR expression in the dorsal striatum. We observed that p-mTOR/mTOR ratio was significantly increased by 139.3 % in 6-OHDA rats compared to sham animals (Fig. 2B; main effect of lesion: F_1,44_ = 3.76, p < 0.05). MSDC-0160 treatment in 6-OHDA rats prevented this increase, but had no significant effect in the sham group (Fig. 2A and 2B; Significant interaction between lesion and treatment: F_1,44_ = 4.76, p < 0.03).

**Fig. 2:**
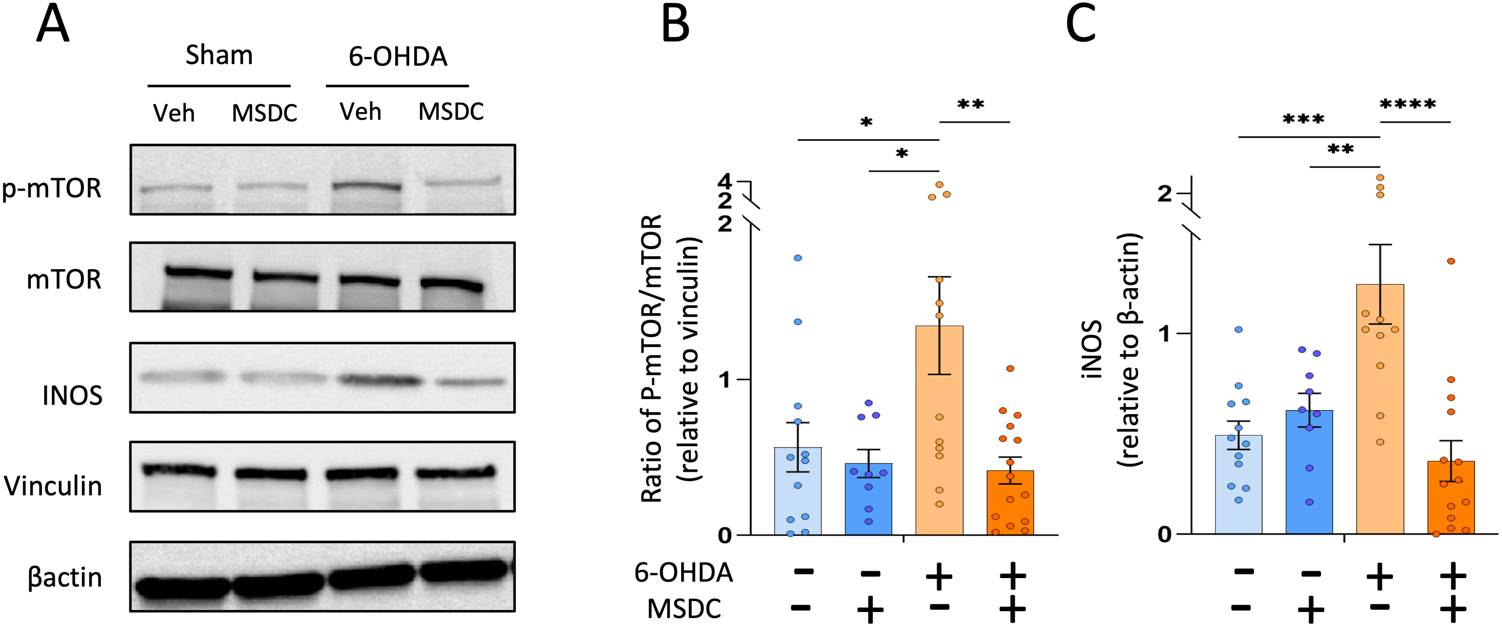
Chronic MSDC-0160 treatment normalized mTOR activity and iNOS expression. (A) Representative western blots illustrating the expression of p-mTOR (Ser^2448^), mTOR, iNOS, vinculin and β-actin in the dorsal striatum of sham and 6-OHDA rats with or without MSDC-0160. (B) Bar graphs showing mean western blot p-mTOR/mTOR ratios relative to vinculin in the dorsal striatum. (C) Bar graphs showing mean western blot iNOS levels relative to β-actin in the dorsal striatum. Mean ± SEM, Two-way ANOVA followed by Tukey’s post-hoc test and correction for multiple comparisons *: p ≤ 0.05, **: p ≤ 0.01, ***: p ≤ 0.001, ****: p ≤ 0.0001

Because neuroinflammatory processes occur in 6-OHDA-induced neuronal damage in the rat [36, 45], we also investigated whether MSDC-0160 administration could modulate iNOS expression in dorsal striatum. We found that MSDC-0160 treatment had no effect in sham animals (Fig. 2A and 2C; post-hoc test of Tukey adjusted p value > 0.91), but completely circumvented the increase in iNOS level induced by 6-OHDA (Fig.2A and 2C; main effect of treatment and significant interaction between lesion and treatment: F_S_ > 7.4, p_S_ < 0.009).

### Chronic MSDC-0160 treatment led to metabolic adaptation of glycolysis and the TCA cycle

We further investigated the potential influence of MSDC-0160 on metabolic alterations induced by 6-OHDA lesion. We focused our analysis on key metabolites related to pyruvate metabolism i.e. glucose, pyruvate and citrate[46]. The expression of the key mitochondrial enzyme pyruvate dehydrogenase (PDH), which links glycolysis to the TCA cycle by generating acetyl-coA from pyruvate[20], was also measured in the dorsal striatum and expressed as a ratio of the inactive phosphorylated form (p-PDH) to total PDH (Fig. 3).

**Fig. 3:**
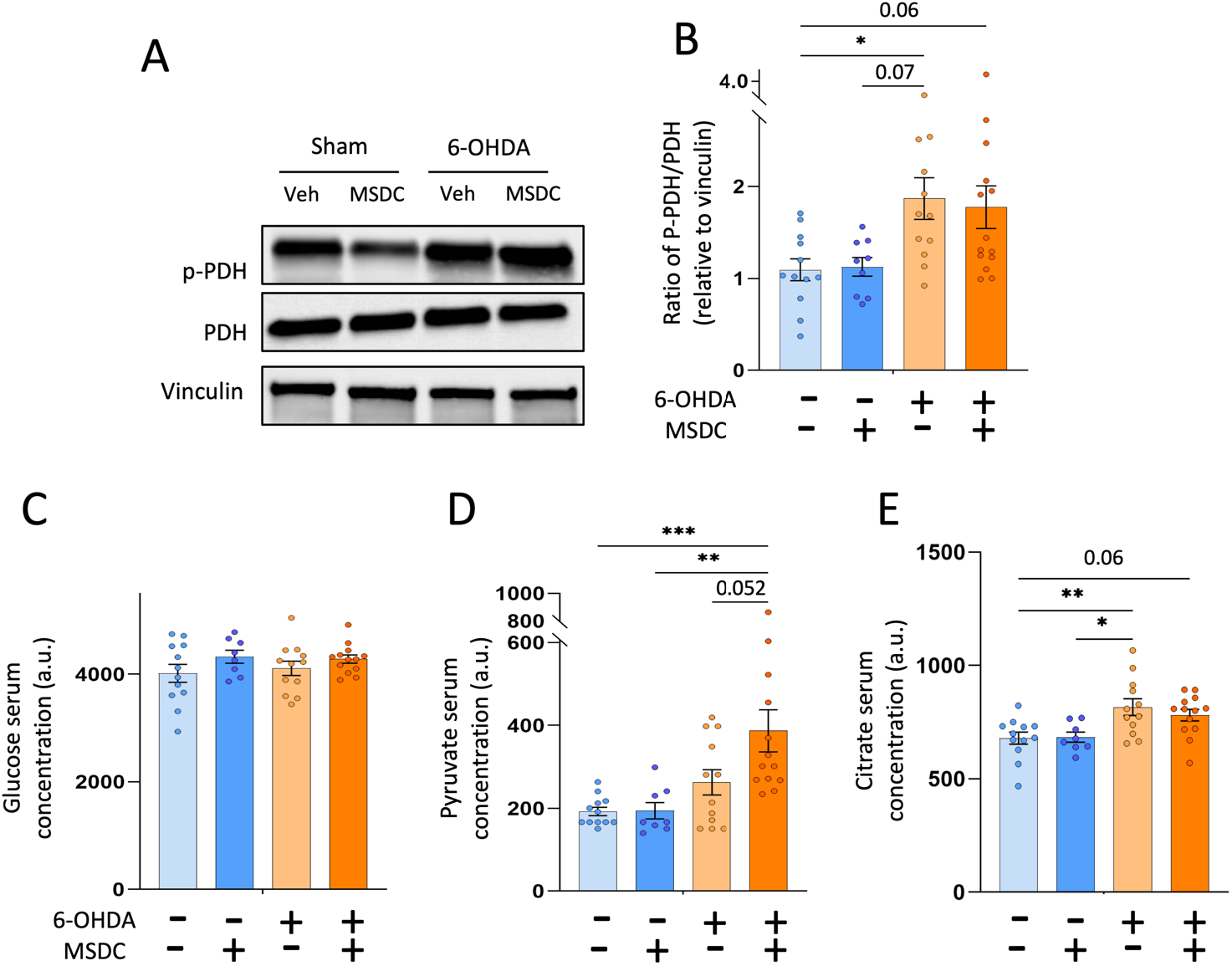
Chronic MSDC-0160 treatment led to metabolic adaptation of glycolysis and the TCA cycle. (A) Representative western blots illustrating the expression of p-PDH (Ser^293^), PDH and vinculin in the dorsal striatum of sham and 6-OHDA rats, with or without MSDC-0160. (B) Bar graphs showing mean western blot p-PDH/PDH ratio relative to vinculin in dorsal striatum. (C-E) Relative serum levels of glucose (C), pyruvate (D) and citrate (E). Mean ± SEM, Two-way ANOVA followed by Tukey’s post-hoc test and correction for multiple comparisons *: p ≤ 0.05, **: p ≤ 0.01, ***: p ≤ 0.001

Neither the injection of 6-OHDA, nor MSDC-0160 treatment altered the circulating level of glucose (Fig. 3C; p = 0.32). Pyruvate levels were slightly increased in lesioned rats and further increased in 6-OHDA animals that received MSDC-0160 treatment compared to control (Fig. 3D; main effect of lesion F_1,44_ = 13.72, p < 0.001, marginal effect of treatment F_1,44_ = 3.15, p = 0.08 and marginal interaction between both factors: F_1,44_ = 2.98, p = 0.09). Finally, we report a treatment-independent increase in the circulating level of citrate in 6-OHDA animals (Fig.3E, main effect of lesion F_1,44_ = 16.72, p < 0.001, but not interaction between lesion and treatment F_1,44_ = 0.37 p = 0.55).

MSDC-0160 treatment alone did not modify the p-PDH/PDH ratio. While a significant 71.5% increase of this ratio was observed in 6-OHDA rats compared to controls, the ratio was not affected by MSDC-0160 treatment (Fig. 3B; main effect of lesion F_1,44_ = 7.41 p = 0.009 but not of treatment F_1,44_ = 0.0003 p = 0.99).

### Chronic MSDC-0160 treatment modified brain energy supply in 6-OHDA rats: Ketogenesis, beta-oxidation and glutamate oxidation

Our previous study[19], reinforced by the present metabolic analysis, suggested decoupling of glycolysis from the TCA cycle, and mitochondrial use of alternative sources of energy[19]. We therefore investigated 3 alternative pathways which can be activated to supply brain energy, i.e. ketogenesis, beta oxidation and glutamate oxidation, by measuring the key enzymes involved in their use, namely, acetyl-CoA acetyltransferase 1 (ACAT1)[47], carnitine palmitoyl transferase 1 (CPT1)[48] and glutamate dehydrogenase (GDH) respectively (Fig. 4E).

**Fig. 4:**
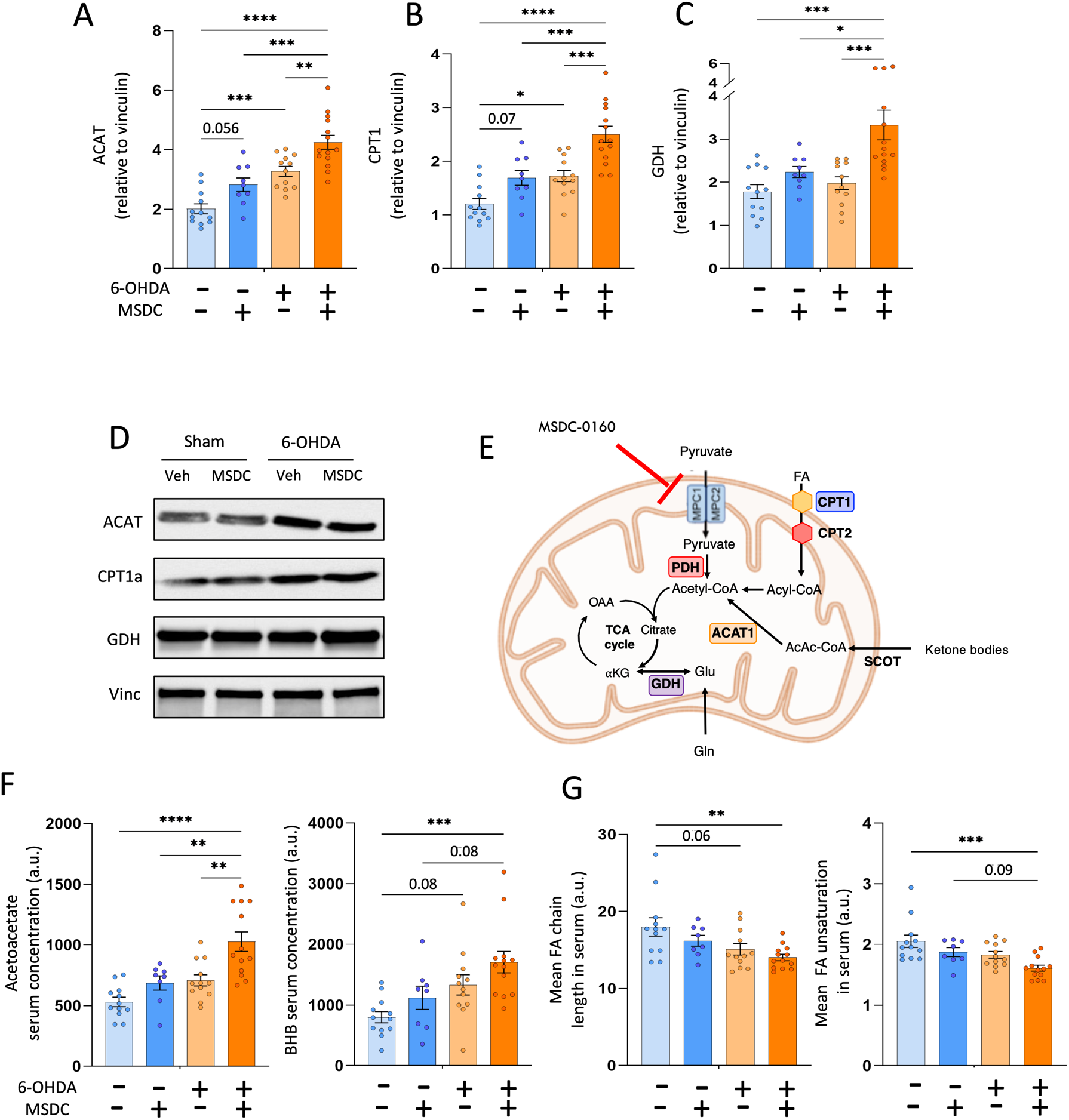
Chronic MSDC-016 treatment modified brain energy supply in 6-OHDA rats: Ketogenesis, beta-oxidation and glutamate oxidation. (A-C) Bar graphs showing mean western blots for ACAT (A), CPT1 (B) and GDH (C), relative to vinculin in the dorsal striatum. (D) Representative western blots illustrating the expression of ACAT, CPT1 and GDH in the dorsal striatum. (E) Schematic diagram summarizing alternative energy sources pathways and their principal actors potentially enhanced by MPC blockage. (F) Relative amplitude of serum ketone body metabolites usable as energetic fuel. (G) Fatty acid chain length and unsaturation, associated with beta oxidation. Mean ± SEM, Two-way ANOVA followed by Tukey’s post-hoc test and correction for multiple comparisons *: p ≤ 0.05, **: p ≤ 0.01, ***: p ≤ 0.001, ****: p ≤ 0.0001. *BHB: Beta-hydroxybutyrate*, *Gln: Glutamine*, *Glu: Glutamate*, *AcAc-Coa: acetoacetyl Coenzyme A*, *FA: Fatty acid*, *OAA: oxaloacetate*, *α-KG: alpha-Ketoglutarate*. *SCOT: 3-ketoacid Coenzyme A transferase*.

First, dorso-striatal expression of ACAT1 significantly increased in 6-OHDA animals compared to shams (Fig. 4A; main effect of lesion F_1,44_ = 42.10, p < 0.001). Moreover, MSDC-0160 treatment increased ACAT1 expression in both sham and 6-OHDA animals (Fig. 4A; main effect of treatment F_1,44_ = 18.54, p < 0.001, without interaction F_1,44_ = 0.15, p = 0.70). Concomitantly, the serum levels of the two main ketone bodies, acetoacetate and β-hydroxybutyrate, rose significantly in 6-OHDA animals and were further increased by treatment with MSDC-0160 (Fig.4F; main effect of lesion and treatment F_S_ > 4.58, p_S_ < 0.03). Similarly, the expression of CPT1, which catalyzes the rate-limiting step of β-oxidation[48], significantly rose with 6-OHDA and MSDC-0160 treatment (Fig. 4B, main effect of lesion and treatment F_S_ > 23.32, p_S_ < 0.001). In parallel, a significant decrease of chain length and unsaturation of fatty acids (FA) was observed in 6-OHDA animals compared to shams (Fig. 4G, main effect of lesion F_S_ > 9.39, p_S_ < 0.003 and marginal effect of treatment F_S_ > 2.83, p_S_ < 0.09).

Finally, 6-OHDA animals treated with MSDC-0160 presented a significant increase of glutamate dehydrogenase (GDH), allowing the entry and use of glutamate in the TCA cycle (Fig. 4C, main effect of lesion and treatment F_S_ > 7.01, p_S_ < 0.01 and marginal interaction F_1,44_ = 3.83; p = 0.07).

## Discussion

There is a growing body of evidence showing that targeting the MPC could constitute a promising approach to find a cure for PD [14, 25, 49]. In the present study, we expanded previous findings regarding the potential neuroprotective effect of MPC inhibition[25, 35] to unilateral 6-OHDA rats. This model, which replicates most of the cell processes encountered in PD[42, 50], represents a phenotypically consistent and standard model of PD. We observed that chronic MSDC-0160 treatment protects against 6-OHDA-induced SNc denervation and associated PD-like motor impairments. Additionally, we found that MSDC-0160 normalized pro-inflammatory and mTOR pathway actors, both impacted by 6-OHDA and implicated in PD physiopathology. Finally, we highlighted multiple actions of MSDC-0160 treatment on mitochondria-related metabolic pathways, promoting the use of alternative energy substrates to glycolysis, in particular by accentuating possible compensatory mechanisms such as ketogenesis that are normally overridden in PD.

Collectively, these results improve our understanding of MSDC-0160 neuroprotective mechanisms and provide the basis for further evaluation of how metabolism reprogramming might impact the course of PD.

In the present study, as expected, unilateral injection of 6-OHDA resulted in damage to the nigrostriatal pathway with significant neurodegeneration in the SNc resulting in denervation of the striatum[42], and leading to a significant decrease of contralateral forelimb use, which generally happens after denervation of at least 70 % of the nigrostriatal pathway[51]. Chronic MSDC-0160 treatment in the 6-OHDA group partially prevented SNc neurodegeneration and was sufficient to almost completely protect against motor impairment. These findings extend results from *C.elegans* and mouse models of PD[25, 49] to a PD rat model with another neurotoxin and therefore further highlight the potential neuroprotective effect of MSDC-0160.

We have also shown that the neuroprotective effect of MSDC-0160 was associated with normalization of mTOR activity in 6-OHDA animals. As a key regulator of cell metabolism and survival, the mTOR signaling pathway, playing a critical role in autophagy, has been found dysregulated in PD[52, 53] and investigated as a potential therapeutic target[4, 54]. Appealingly, TZDs can modulate mTOR activity, particularly the mTORC1 complex involved in autophagy [55, 56]. Moreover, it has been shown that autophagic signals could modulate neuroinflammation[57–59], which is instrumental in neuronal cell death and particularly involved in the progression of neurodegeneration[9, 60]. Furthermore, neuroinflammatory processes in PD, induced by released neurotoxic factors such as nitric oxide, are mediated by iNOS[61, 62]. Well in line with previous studies reporting increase of nitrite levels[61, 63] in 6-OHDA animals, we observed a stronger expression of iNOS in untreated lesioned animals, which was alleviated by MSDC-0160. Consistently, NOS inhibitors have been investigated as a potential therapeutic strategy in 6-OHDA animals and are shown to protect DA neurons of the nigrostriatal pathway and reduce the associated motor deficits[61, 64, 65]. Overall, these results strongly suggest that the MSDC-0160 neuroprotective effect observed in our study may be achieved through a mTOR modulation and an associated reduction of neuroinflammation.

In addition to these mechanisms, global energy failure linked to mitochondrial dysfunction represents an hallmark of neuronal death in PD[18, 66], and markers of these dysfunctions, such as inhibition of complex I, have been reported in brain and blood of PD patients[3]. Under normal physiological conditions, glucose is the dominant exogenous energy substrate in the brain[46]. It is converted into pyruvate, the master fuel for the TCA cycle in mitochondria[15]. In the present study, neither 6-OHDA nor MDSC-0160 altered blood glucose levels, but blood pyruvate increased in both 6-OHDA groups. This pyruvate accumulation is consistent with the increased p-PDH/PDH ratio observed in the lesioned rats since the activity of PDH regulates entry of pyruvate into the TCA cycle. Abnormally increased levels of pyruvate and decreased levels of PDH have already been observed in PD models and patients[67, 68]. Moreover, in rodents, downregulation of PDH was associated with dopaminergic degeneration and could result in motor impairments similar to those observed in other PD animal models[69]. The neuroprotective effect of MSDC-0160 was not associated with a reduction of PDH expression compared to untreated 6-OHDA animals which is consistent with previous studies reporting that TZDs do not influence PDH activation[70] and that MPC knock-down leads to metabolic reprogramming distinct from PDH inhibition[24]. Surprisingly, despite diminution of PDH in 6-OHDA animals, limiting the use of pyruvate as fuel[71], citrate, the first metabolite of the TCA cycle, was in fact increased in these animals, suggesting its potential supply form one or several alternative sources[72].

Among them, ketone bodies represent one of the main alternative fuels for the brain (see fig 4E). Produced by oxidation of fatty acid in the liver, ketones bodies, such as beta-hydroxybutyrate (BHB) and acetoacetate, are transported by blood and are able to cross the blood–brain barrier to produce acetyl-CoA thus supplying the TCA cycle. In the present study, we have shown that MSDC-0160 treatment increased the blood levels of BHB and acetoacetate in sham animals, and even further increased them in lesioned animals. These observations were reinforced by MSDC-0160-dependent upregulation of ACAT1, the pivotal enzyme of ketogenesis[47, 73]. Interestingly, increased ketogenesis has been previously studied in the PD context and, as in our experiments, has been shown to alleviate motor dysfunction, reduce neuronal loss in the SNc and down-regulate neuroinflammation[74, 75]. Moreover, recent studies highlighted crucial roles for ketone bodies in cell metabolism, in particular in the maintenance of energy production in condition such as MPC inhibition[24, 27]. Furthermore, a ketogenic diet has been shown to prevent the lethality induced by MPC knock-out in mice[23]. Finally, substitution of glucose by ketone bodies inactivates PDH[76–78] and might also indirectly affect pyruvate utilization by competing for entry into mitochondria, which could explain our result associated with pyruvate metabolism[72, 79].

Concomitantly, the decrease of mean chain length and mean unsaturation of fatty acids that we observed in blood of 6-OHDA animals treated with MSDC-0160 suggests an increase of β-oxidation[80], consistent with the increase of ketone bodies. These results are supported by brain expression of CPT1, catalyzing the rate-limiting step of β-oxidation[48], which appeared increased by 6-OHDA and even more by MSDC-0160 treatment. Appealingly, CPT1 can be modulated by TZDs[56]. Moreover, variation in fatty acid chain lengths, possibly maintaining acetyl-CoA synthesis and therefore energy homeostasis, have also been observed in other animal models of PD[81]. Finally, CPT1 can promote human mesenchymal stem cell survival under glucose deprivation through an enhanced availability of fatty acids as fuel substrates for ATP generation[82].

In addition to these alternative sources, we have shown that MSDC-0160 increased the expression of glutamate dehydrogenase (GDH), which was even further increased in the 6-OHDA-lesioned rats, supporting the possibility of glutamate utilization for cell metabolism under MPC inhibition. Indeed, GDH allows glutamate to supply the TCA cycle by the formation of α-ketoglutarate. When the glucose supply is sufficient, and energy status adequate to cell needs, this pathway is less active. However, when energy demand is increased, as in the context of PD pathology, GDH activity is increased[83] in order to support the anaplerotic needs of the cell[27, 49]. Moreover, this switching to glutamate oxidation may reduce the glutamate pool and therefore excitotoxic neuronal death[27, 49, 84], contributing to the neuroprotective effects of MPC inhibition.

In summary, our results reveal that in unilateral PD rats, the neuroprotective effect of MSDC-0160 treatment includes several distinct mechanisms which appear to be upstream of metabolic reprogramming. Indeed, as already described, MSDC-0160 treatment can correct mechanisms related to neuronal death such as autophagy and neuroinflammation. Moreover, our results strongly suggest that the effect of MSDC-0160 treatment could be seen as a reprogramming of energy production inside the brain. Indeed, we demonstrated that MSDC-0160 treatment increased the use of alternative substrates to supply the energy needs in the PD context and to try to maintain energy homeostasis[84–86]. Compared with targeting a single pathway, such as the ketogenic diet, which improves PD symptomatology[78, 87] but seems difficult to foresee in the long term[88], this multi-mechanism action, leading to global reorganization of cell metabolism, appears preferable. Notably, the present study highlights that MPC can be targeted by clinically safe drugs such as MSDC-0160 and shows promising effects that could provide a useful treatment for PD.

## Conclusion

This study contributes to a clearer understanding of the potential of MSDC-0160 as a neuroprotective treatment, by reprogramming metabolism through inhibition of the MPC. Although much remains to be learned about the underlying mechanisms, the translational potential of MPC inhibition for the treatment of PD is strongly supported by these findings.

## Abbreviations

6-OHDA: 6-hydroxydopamine
DS: Dorsal striatum
mTOR: Mammalian target of rapamycin
MPC: Mitochondrial pyruvate carrier
MPTP: 1-methyl-4-phenyl-1,2,3,6-tetrahydropyridine
NMR: Nuclear magnetic resonance
PD: Parkinson’s disease
SNc: Substantia nigra pars compacta
TCA: Tricarboxylic acid
TH-IR: Tyrosine hydroxylase immunoreactivity
TZD: thiazolidinedione

## Supplementary Information

**Additional file 1. Fig. S1:** Location of cannula implantations within the SNc in the different groups.

## Acknowledgement

The authors would like to thank Adrien Favier and Alicia Vallet from IBS (Grenoble) for technical assistance concerning NMR experiments and IR-RMN-THC Fr3050 CNRS for financial support for conducting the research. They also thank Magali Bartolomucci for her technical help regarding cylinder test experiment.

## Declaration

### Author contributions

Research: D.M, R.G, J.C, S.C, E.L.B, S.B, F.F

Conducted experiments: D.M, R.G

Designed research: D.M, S.B, F.F, J.C

Analysed data D.M, S.B, F.F

Wrote the manuscript D.M, S.B, F.F with the help of the other authors

### Availability of data and materials

All data generated or analyzed during this study are included in this article and supplementary information files.

### Ethic approval and consent to participate

All protocols complied with the European Union 2010 Animal Welfare Act and the French directive 2010/63 and were approved by the French national ethics committee (2013/113) n° 004 and by local ethical committee C2EA84.

### Consent for publication

Not applicable

## References

1. Poewe W, Seppi K, Tanner CM, et al (2017) Parkinson disease. Nature Reviews Disease Primers. https://doi.org/10.1038/nrdp.2017.13

2. Nicoletti V, Palermo G, Del Prete E, et al (2021) Understanding the Multiple Role of Mitochondria in Parkinson’s Disease and Related Disorders: Lesson From Genetics and Protein–Interaction Network. Frontiers in Cell and Developmental Biology

3. Camilleri A, Vassallo N (2014) The Centrality of mitochondria in the pathogenesis and treatment of Parkinson’s disease. CNS Neuroscience and Therapeutics

4. Bové J, Martínez-Vicente M, Vila M (2011) Fighting neurodegeneration with rapamycin: Mechanistic insights. Nature Reviews Neuroscience

5. Tansey MG, Goldberg MS (2010) Neuroinflammation in Parkinson’s disease: Its role in neuronal death and implications for therapeutic intervention. Neurobiology of Disease

6. Roca-Agujetas V, De Dios C, Lestón L, et al (2019) Recent Insights into the Mitochondrial Role in Autophagy and Its Regulation by Oxidative Stress. Oxidative Medicine and Cellular Longevity

7. Zheng X, Boyer L, Jin M, et al (2016) Alleviation of neuronal energy deficiency by mtor inhibition as a treatment for mitochondria-related neurodegeneration. eLife. https://doi.org/10.7554/eLife.13378

8. Brown GC, Bal-Price A (2003) Inflammatory Neurodegeneration Mediated by Nitric Oxide, Glutamate, and Mitochondria. Molecular Neurobiology

9. Di Filippo M, Chiasserini D, Tozzi A, et al (2010) Mitochondria and the link between neuroinflammation and neurodegeneration. Journal of Alzheimer’s Disease

10. Athauda D, Foltynie T (2015) The ongoing pursuit of neuroprotective therapies in Parkinson disease. Nature Reviews Neurology

11. Kalia L V., Kalia SK, Lang AE (2015) Disease-modifying strategies for Parkinson’s disease. Movement Disorders

12. Olanow CW, Kieburtz K, Schapira AH v (2008) Why have we failed to achieve neuroprotection in Parkinson’s disease? Annals of Neurology 64

13. Rey F, Ottolenghi S, Giallongo T, et al (2021) Mitochondrial metabolism as target of the neuroprotective role of erythropoietin in Parkinson’s disease. Antioxidants. https://doi.org/10.3390/antiox10010121

14. Quansah E, Peelaerts W, Langston JW, et al (2018) Targeting energy metabolism via the mitochondrial pyruvate carrier as a novel approach to attenuate neurodegeneration. Molecular Neurodegeneration

15. Gray LR, Tompkins SC, Taylor EB (2014) Regulation of pyruvate metabolism and human disease. Cellular and Molecular Life Sciences

16. Martínez-Reyes I, Chandel NS (2020) Mitochondrial TCA cycle metabolites control physiology and disease. Nature Communications

17. Ahmed SS, Santosh W, Kumar S, Christlet HTT (2009) Metabolic profiling of Parkinson’s disease: Evidence of biomarker from gene expression analysis and rapid neural network detection. Journal of Biomedical Science. https://doi.org/10.1186/1423-0127-16-63

18. Anandhan A, Jacome MS, Lei S, et al (2017) Metabolic Dysfunction in Parkinson’s Disease: Bioenergetics, Redox Homeostasis and Central Carbon Metabolism. Brain Research Bulletin

19. Mallet D, Dufourd T, Decourt M, et al (2021) Early diagnosis of Parkinson’s disease: A cross-species biomarker. bioRxiv 2021.10.04.462993. https://doi.org/10.1101/2021.10.04.462993

20. Zangari J, Petrelli F, Maillot B, Martinou JC (2020) The multifaceted pyruvate metabolism: Role of the mitochondrial pyruvate carrier. Biomolecules. https://doi.org/10.3390/biom10071068

21. Buchanan JL, Taylor EB (2020) Mitochondrial pyruvate carrier function in health and disease across the lifespan. Biomolecules

22. Bender T, Martinou JC (2016) The mitochondrial pyruvate carrier in health and disease: To carry or not to carry? Biochimica et Biophysica Acta - Molecular Cell Research

23. Vanderperre B, Herzig S, Krznar P, et al (2016) Embryonic Lethality of Mitochondrial Pyruvate Carrier 1 Deficient Mouse Can Be Rescued by a Ketogenic Diet. PLoS Genetics. https://doi.org/10.1371/journal.pgen.1006056

24. Vacanti NM, Divakaruni AS, Green CR, et al (2014) Regulation of substrate utilization by the mitochondrial pyruvate carrier. Molecular Cell 56:425–435. https://doi.org/10.1016/j.molcel.2014.09.024

25. Ghosh A, Tyson T, George S, et al (2016) Mitochondrial pyruvate carrier regulates autophagy, inflammation, and neurodegeneration in experimental models of Parkinson’s disease. Science Translational Medicine 8:. https://doi.org/10.1126/scitranslmed.aag2210

26. Divakaruni AS, Wiley SE, Rogers GW, et al (2013) Thiazolidinediones are acute, specific inhibitors of the mitochondrial pyruvate carrier. Proceedings of the National Academy of Sciences of the United States of America. https://doi.org/10.1073/pnas.1303360110

27. Divakaruni AS, Wallace M, Buren C, et al (2017) Inhibition of the mitochondrial pyruvate carrier protects from excitotoxic neuronal death. Journal of Cell Biology. https://doi.org/10.1083/jcb.201612067

28. Brauer R, Bhaskaran K, Chaturvedi N, et al (2015) Glitazone treatment and incidence of parkinson’s disease among people with diabetes: A retrospective cohort study. PLoS Medicine. https://doi.org/10.1371/journal.pmed.1001854

29. Hernandez-Baltazar D, Zavala-Flores LM, Villanueva-Olivo A (2017) The 6-hydroxydopamine model and parkinsonian pathophysiology: Novel findings in an older model. Neurología (English Edition). https://doi.org/10.1016/j.nrleng.2015.06.019

30. Wachter B, Schürger S, Rolinger J, et al (2010) Effect of 6-hydroxydopamine (6-OHDA) on proliferation of glial cells in the rat cortex and striatum: Evidence for de-differentiation of resident astrocytes. Cell and Tissue Research. https://doi.org/10.1007/s00441-010-1061-x

31. Haas SJP, Zhou X, Machado V, et al (2016) Expression of tgfβ1 and inflammatory markers in the 6-hydroxydopamine mouse model of Parkinson’s disease. Frontiers in Molecular Neuroscience. https://doi.org/10.3389/fnmol.2016.00007

32. Drui G, Carnicella S, Carcenac C, et al (2014) Loss of dopaminergic nigrostriatal neurons accounts for the motivational and affective deficits in Parkinson’s disease. Molecular psychiatry 19:358–67. https://doi.org/10.1038/mp.2013.3

33. Favier M, Duran T, Carcenac C, et al (2014) Pramipexole reverses Parkinson’s disease-related motivational deficits in rats. Movement Disorders 29:912–920. https://doi.org/10.1002/mds.25837

34. Paxinos G, Watson C (1997) The Rat Brain in Stereotaxic Coordinates. Academic Press, San Diego 3rd:

35. Peelaerts W, Bergkvist L, George S, et al (2020) Inhibiting the mitochondrial pyruvate carrier does not ameliorate synucleinopathy in the absence of inflammation or metabolic deficits. bioRxiv

36. Rentsch P, Stayte S, Morris GP, Vissel B (2019) Time dependent degeneration of the nigrostriatal tract in mice with 6-OHDA lesioned medial forebrain bundle and the effect of activin A on l-Dopa induced dyskinesia. BMC Neuroscience. https://doi.org/10.1186/s12868-019-0487-7

37. Magno LA, Collodetti M, Tenza-Ferrer H, Romano-Silva M (2019) Cylinder Test to Assess Sensory-Motor Function in a Mouse Model of Parkinson’s Disease. BIO-PROTOCOL. https://doi.org/10.21769/bioprotoc.3337

38. Butler TAJ, Paul JW, Chan EC, et al (2019) Misleading westerns: Common quantification mistakes in western blot densitometry and proposed corrective measures. BioMed Research International. https://doi.org/10.1155/2019/5214821

39. Beckonert O, Keun HC, Ebbels TMD, et al (2007) Metabolic profiling, metabolomic and metabonomic procedures for NMR spectroscopy of urine, plasma, serum and tissue extracts. Nat Protoc 2:2692–2703. https://doi.org/10.1038/nprot.2007.376

40. Wishart DS, Feunang YD, Marcu A, et al (2018) HMDB 4.0: The human metabolome database for 2018. Nucleic Acids Research. https://doi.org/10.1093/nar/gkx1089

41. Zancanaro C, Nano R, Marchioro C, et al (1994) Magnetic resonance spectroscopy investigations of brown adipose tissue and isolated brown adipocytes. Journal of Lipid Research. https://doi.org/10.1016/s0022-2275(20)39925-9

42. Magnard R, Vachez Y, Carcenac C, et al (2016) What can rodent models tell us about apathy and associated neuropsychiatric symptoms in Parkinson’s disease? Transl Psychiatry 6:e753. https://doi.org/10.1038/tp.2016.17

43. Schwarting RKW, Huston JP (1996) Unilateral 6-hydroxydopamine lesions of meso-striatal dopamine neurons and their physiological sequelae. Progress in Neurobiology

44. He X, Yuan W, Li Z, Feng J (2017) An autophagic mechanism is involved in the 6-hydroxydopamine-induced neurotoxicity in vivo. Toxicology Letters 280:29–40. https://doi.org/10.1016/j.toxlet.2017.08.006

45. Walsh S, Finn DP, Dowd E (2011) Time-course of nigrostriatal neurodegeneration and neuroinflammation in the 6-hydroxydopamine-induced axonal and terminal lesion models of Parkinson’s disease in the rat. Neuroscience. https://doi.org/10.1016/j.neuroscience.2010.12.005

46. Dienel GA (2019) Brain glucose metabolism: Integration of energetics with function. Physiological Reviews. https://doi.org/10.1152/physrev.00062.2017

47. Abdelkreem E, Harijan RK, Yamaguchi S, et al (2019) Mutation update on ACAT1 variants associated with mitochondrial acetoacetyl-CoA thiolase (T2) deficiency. Human Mutation. https://doi.org/10.1002/humu.23831

48. Wolfgang MJ, Kurama T, Dai Y, et al (2006) The brain-specific carnitine palmitoyltransferase-1c regulates energy homeostasis. Proceedings of the National Academy of Sciences of the United States of America. https://doi.org/10.1073/pnas.0602205103

49. Tang BL (2019) Targeting the mitochondrial pyruvate carrier for neuroprotection. Brain Sciences

50. Hernandez-Baltazar D, Nadella R, Rovirosa-Hernandez M de J, et al (2018) Animal Model of Parkinson Disease: Neuroinflammation and Apoptosis in the 6-Hydroxydopamine-Induced Model. In: Experimental Animal Models of Human Diseases - An Effective Therapeutic Strategy

51. Schwarting RKW, Huston JP (1996) The unilateral 6-hydroxydopamine lesion model in behavioral brain research. Analysis of functional deficits, recovery and treatments. Progress in Neurobiology

52. Malagelada C, Ryu EJ, Biswas SC, et al (2006) RTP801 is elevated in Parkinson brain substantia nigral neurons and mediates death in cellular models of Parkinson’s disease by a mechanism involving mammalian target of rapamycin inactivation. Journal of Neuroscience. https://doi.org/10.1523/JNEUROSCI.3292-06.2006

53. Lan A ping, Chen J, Zhao Y, et al (2017) mTOR Signaling in Parkinson’s Disease. NeuroMolecular Medicine

54. Zhu Z, Yang C, Iyaswamy A, et al (2019) Balancing mTOR signaling and autophagy in the treatment of Parkinson’s disease. International Journal of Molecular Sciences

55. Colca JR (2015) The TZD insulin sensitizer clue provides a new route into diabetes drug discovery. Expert Opinion on Drug Discovery

56. Colca JR, Mcdonald WG, Kletzien RF (2014) Mitochondrial target of thiazolidinediones. Diabetes, Obesity and Metabolism

57. Palavra F, Ambrósio AF, Reis F (2016) mTOR and Neuroinflammation. In: Molecules to Medicine with mTOR: Translating Critical Pathways into Novel Therapeutic Strategies

58. Zheng HF, Yang YP, Hu LF, et al (2013) Autophagic Impairment Contributes to Systemic Inflammation-Induced Dopaminergic Neuron Loss in the Midbrain. PLoS ONE. https://doi.org/10.1371/journal.pone.0070472

59. Cho KS, Lee JH, Cho J, et al (2018) Autophagy Modulators and Neuroinflammation. Current Medicinal Chemistry. https://doi.org/10.2174/0929867325666181031144605

60. Hirsch EC, Vyas S, Hunot S (2012) Neuroinflammation in Parkinson’s disease. Parkinsonism and Related Disorders. https://doi.org/10.1016/s1353-8020(11)70065-7

61. Singh S, Kumar S, Dikshit M (2010) Involvement of the mitochondrial apoptotic pathway and nitric oxide synthase in dopaminergic neuronal death induced by 6-hydroxydopamine and lipopolysaccharide. Redox Report. https://doi.org/10.1179/174329210X12650506623447

62. Hunot S, Boissière F, Faucheux B, et al (1996) Nitric oxide synthase and neuronal vulnerability in Parkinson’s disease. Neuroscience. https://doi.org/10.1016/0306-4522(95)00578-1

63. Di Matteo V, Pierucci M, Benigno A, et al (2009) Involvement of nitric oxide in nigrostriatal dopaminergic system degeneration: A neurochemical study. In: Annals of the New York Academy of Sciences

64. Singh S, Das T, Ravindran A, et al (2005) Involvement of nitric oxide in neurodegeneration: A study on the experimental models of Parkinson’s disease. Redox Report. https://doi.org/10.1179/135100005X38842

65. Li M, Dai FR, Du XP, et al (2012) Neuroprotection by silencing iNOS expression in a 6-OHDA model of Parkinson’s disease. Journal of Molecular Neuroscience. https://doi.org/10.1007/s12031-012-9814-5

66. Pathak D, Berthet A, Nakamura K (2013) Energy failure: Does it contribute to neurodegeneration? Annals of Neurology. https://doi.org/10.1002/ana.24014

67. Ahmed SSSJ, Santosh W, Kumar S, Christlet HTT (2009) Metabolic profiling of Parkinson’s disease: evidence of biomarker from gene expression analysis and rapid neural network detection. JOURNAL OF BIOMEDICAL SCIENCE 16:. https://doi.org/10.1186/1423-0127-16-63

68. Henchcliffe C, Shungu DC, Mao X, et al (2008) Multinuclear magnetic resonance spectroscopy for in vivo assessment of mitochondrial dysfunction in Parkinson’s disease. In: Annals of the New York Academy of Sciences

69. Ojano-Dirain C, Glushakova LG, Zhong L, et al (2010) An animal model of PDH deficiency using AAV8-siRNA vector-mediated knockdown of pyruvate dehydrogenase E1α. Molecular Genetics and Metabolism 101:183–191. https://doi.org/10.1016/j.ymgme.2010.07.008

70. Shannon CE, Daniele G, Galindo C, et al (2017) Pioglitazone inhibits mitochondrial pyruvate metabolism and glucose production in hepatocytes. FEBS Journal. https://doi.org/10.1111/febs.13992

71. Patel MS, Nemeria NS, Furey W, Jordan F (2014) The pyruvate dehydrogenase complexes: Structure-based function and regulation. Journal of Biological Chemistry

72. Roberts EL (2007) The support of energy metabolism in the central nervous system with substrates other than glucose. In: Handbook of Neurochemistry and Molecular Neurobiology: Brain Energetics. Integration of Molecular and Cellular Processes

73. Rose J, Brian C, Pappa A, et al (2020) Mitochondrial Metabolism in Astrocytes Regulates Brain Bioenergetics, Neurotransmission and Redox Balance. Frontiers in Neuroscience

74. Tieu K, Perier C, Caspersen C, et al (2003) D-β-Hydroxybutyrate rescues mitochondrial respiration and mitigates features of Parkinson disease. Journal of Clinical Investigation. https://doi.org/10.1172/JCI200318797

75. Yang X, Cheng B (2010) Neuroprotective and anti-inflammatory activities of ketogenic diet on MPTP-induced neurotoxicity. Journal of Molecular Neuroscience. https://doi.org/10.1007/s12031-010-9336-y

76. Reed LJ (1981) Regulation of Mammalian Pyruvate Dehydrogenase Complex by a Phosphorylation–Dephosphorylation Cycle. In: Current Topics in Cellular Regulation

77. Patel MS, Korotchkina LG (2003) The biochemistry of the pyruvate dehydrogenase complex. Biochemistry and Molecular Biology Education

78. Puchalska P, Crawford PA (2017) Multi-dimensional Roles of Ketone Bodies in Fuel Metabolism, Signaling, and Therapeutics. Cell Metabolism

79. Booth RFG, Clark JB (1981) Energy Metabolism in Rat Brain: Inhibition of Pyruvate Decarboxylation by 3-Hydroxybutyrate in Neonatal Mitochondria. Journal of Neurochemistry. https://doi.org/10.1111/j.1471-4159.1981.tb05306.x

80. Chance DS, McIntosh MK (1994) Rates of beta-oxidation of fatty acids of various chain lengths and degrees of unsaturation in highly purified peroxisomes isolated from rat liver. Comparative Biochemistry and Physiology -- Part B: Biochemistry and. https://doi.org/10.1016/0305-0491(94)90011-6

81. Worth AJ, Basu SS, Snyder NW, et al (2014) Inhibition of neuronal cell mitochondrial complex i with rotenone increases lipid β-oxidation, supporting acetyl-coenzyme a levels. Journal of Biological Chemistry. https://doi.org/10.1074/jbc.M114.591354

82. Roa-Mansergas X, Fadó R, Atari M, et al (2018) CPT1C promotes human mesenchymal stem cells survival under glucose deprivation through the modulation of autophagy. Scientific Reports. https://doi.org/10.1038/s41598-018-25485-7

83. Zaganas I V., Kanavouras K, Borompokas N, et al (2014) The odyssey of a young gene: Structure-function studies in human glutamate dehydrogenases reveal evolutionary-acquired complex allosteric regulation mechanisms. Neurochemical Research

84. Kim AY, Baik EJ (2019) Glutamate Dehydrogenase as a Neuroprotective Target Against Neurodegeneration. Neurochemical Research. https://doi.org/10.1007/s11064-018-2467-1

85. Greco T, Glenn TC, Hovda DA, Prins ML (2016) Ketogenic diet decreases oxidative stress and improves mitochondrial respiratory complex activity. Journal of Cerebral Blood Flow and Metabolism. https://doi.org/10.1177/0271678X15610584

86. Rojas-Morales P, Pedraza-Chaverri J, Tapia E (2020) Ketone bodies, stress response, and redox homeostasis. Redox Biology

87. Brietzke E, Mansur RB, Subramaniapillai M, et al (2018) Ketogenic diet as a metabolic therapy for mood disorders: Evidence and developments. Neuroscience and Biobehavioral Reviews

88. Włodarek D (2019) Role of ketogenic diets in neurodegenerative diseases (Alzheimer’s disease and parkinson’s disease). Nutrients

