## Supplementary figures and images for "Re-routing metabolism by the mitochondrial pyruvate carrier inhibitor MSDC-0160 attenuates neurodegeneration in a rat model of Parkinson’s disease"

### Additional file 1: Fig. S1

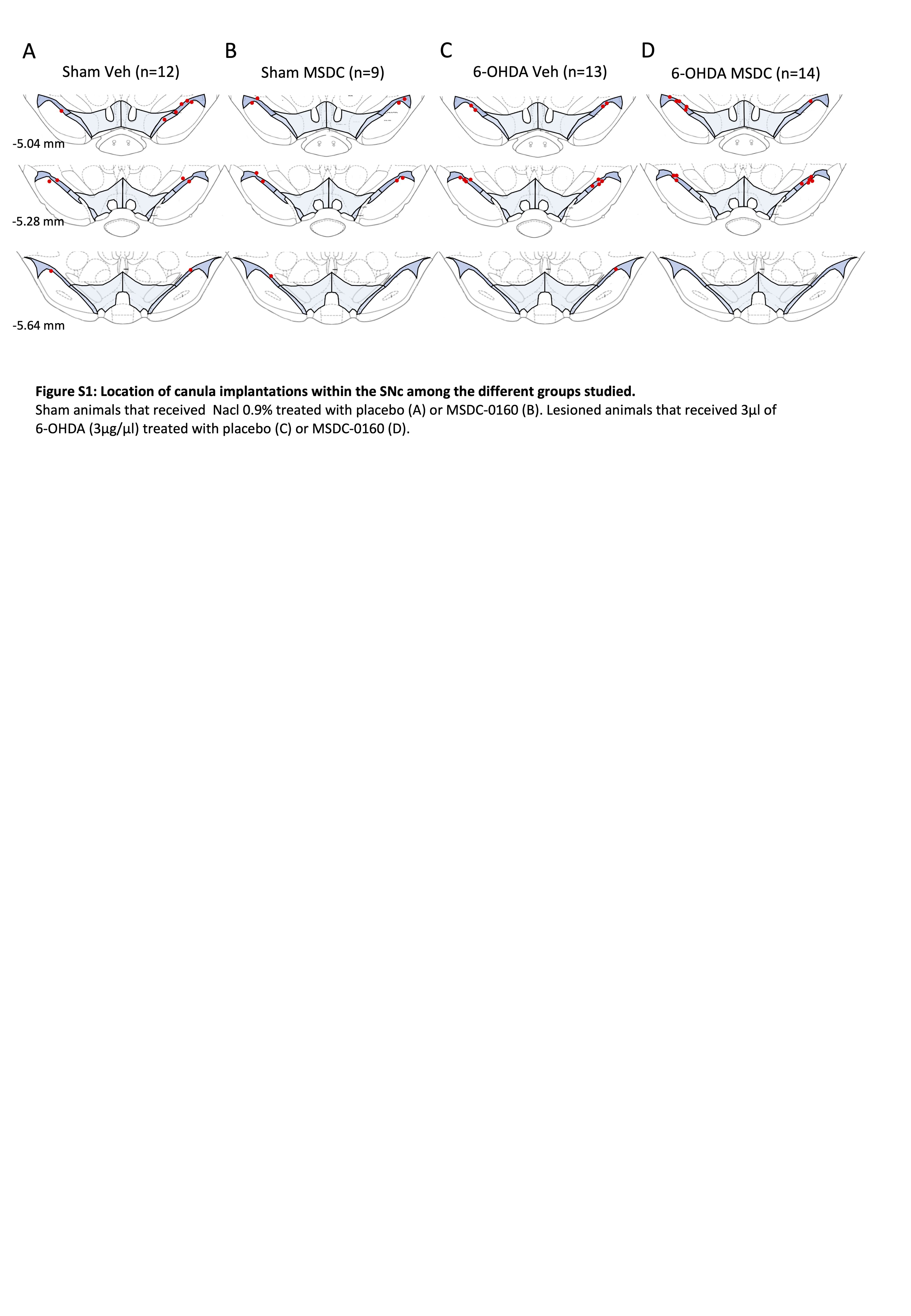
